# Dopamine and serotonin interplay for valence-based spatial learning

**DOI:** 10.1101/2021.03.04.433867

**Authors:** Carlos Wert-Carvajal, Melissa Reneaux, Tatjana Tchumatchenko, Claudia Clopath

**Author notes:** Correspondence (CC), (TT).

## Abstract

Dopamine and serotonin are important modulators of synaptic plasticity and their action has been linked to our ability to learn the positive or negative outcomes or valence learning. In the hippocampus, both neuromodulators affect long-term synaptic plasticity but play different roles in the encoding of uncertainty or predicted reward. Here, we examine the differential role of these modulators on learning speed and cognitive flexibility in a navigational model. We compare two reward-modulated spike time-dependent plasticity (R-STDP) learning rules to describe the action of these neuromodulators. Our results show that the interplay of dopamine (DA) and serotonin (5-HT) improves overall learning performance and can explain experimentally reported differences in spatial task performance. Furthermore, this system allows us to make predictions regarding spatial reversal learning.

## 1 Introduction

The interplay between dopamine (DA) and serotonin, or 5-hydroxytryptamine (5-HT), regulates cognitive functions underpinning decision-making. However, its behavioral consequences and integration into a control system remain elusive (Barnes and Sharp, 1999; Dayan and Huys, 2015). DA has long been characterized as a plasticity modulator that encodes uncertainty or prediction error in reinforcement learning (Schultz et al., 1997; Sutton and Barto, 2018). Unlike other neuromodulators, such as acetylcholine or noradrenaline (Frémaux and Gerstner, 2015), the role of 5-HT is less clear and it has been hypothesized to contribute to aversive processing, analogous to the function of DA in positive or appetite-driven rewards (Rogers, 2011; Cools et al., 2011; Crockett et al., 2012; Cohen et al., 2015; Fischer and Ullsperger, 2017). 5-HT also regulates traits of the social brain such as depression and impulsiveness (Rogers, 2011; Dalley and Roiser, 2012). In the hippocampus, which is critical for spatial memory formation (O’Keefe and Dostrovsky, 1971; O’Keefe and Nadel, 1978), DA and 5-HT have been studied in the context of valence-based learning (Fischer and Ullsperger, 2017; Fernandez et al., 2017; Schmidt et al., 2017; Waider et al., 2019). Even if true opponency is not well-established (Daw et al., 2002; Boureau and Dayan, 2011), evidence suggests that the antagonistic effects of DA and 5-HT can explain neural activity during reward-driven learning (Crockett et al., 2012; Cohen et al., 2015; Matias et al., 2017). Notably, DA has been shown to induce long-term potentiation (LTP) in reward-guided navigation (Brzosko et al., 2015; Palacios-Filardo and Mellor, 2019), and 5-HT longterm depression (LTD) for some receptor-specific hippocampal areas (Kemp and Manahan-Vaughan, 2004, 2005; Berumen et al., 2012; Wawra et al., 2014; Lecouflet et al., 2020). However, 5-HT could also produce LTP or metaplasticity regulation (Wang and Arvanov, 1998; Hagena and Manahan-Vaughan, 2017; Teixeira et al., 2018).

Motivated by these experimental findings, we present a mathematical model of valence-based learning in the hippocampus which details the antagonistic roles of DA and 5-HT for long-term synaptic plasticity. To this end, we use available biological data describing the dynamics of both neuromodulators and present a stable neoHebbian three-factor learning rule (Frémaux and Gerstner, 2015; Gerstner et al., 2018; Zannone et al., 2018) characterising their effect in synapses. By evaluating and optimizing two spike time-dependent plasticity (STDP) rules during forward learning in a navigational task, we find that 5-HT increases training performance. Finally, we show that the proposed interplay of 5-HT and DA resembles behavioral evidence and can shape the adaptation in reversal learning (Matias et al., 2017).

## 2 Results and discussion

The valence system that we propose for DA and 5-HT contributions highlights the functional importance of the competition between timing-dependent long-term potentiation (t-LTP) and depression (t-LTD) during rewarding and punishing reinforcement cues. We followed a navigational hippocampal model (Foster et al., 2000; Vasilaki et al., 2009; Frémaux et al., 2013; Brzosko et al., 2017) with a feed-forward network of presynaptic place cells and a layer of postsynaptic action neurons (***Figure 1A***; Materials and methods). In reward-modulated spike timing-dependent plasticity (R-STDP) weight change is a function of the firing difference and the action of a neuromodulator (***Figure 1B***). We used previously reported data describing the STDP window for DA in the hippocampus (Brzosko et al., 2015, 2017) and assumed that LTD-inducing effects of 5-HT can be captured by an anti-causal learning window, as shown in cortical 5-HT_2C_ receptors (He et al., 2015; ***Figure 1B***). We found that varying STDP windows for 5-HT and DA produced similar navigational outcomes through an equal decay which we chose for further analysis (***Figure S1***). Temporal discrimination of neural activity leading up to the reinforcement signal was achieved through an eligibility trace (Gerstner et al., 2018), also known as proto-weight, for which there is evidence for DA (Brzosko et al., 2017) and 5-HT (He et al., 2015; ***Figure 1C***). The eligibility trace of 5-HT, adapted from the neocortex, presents slower dynamics than that of DA, in the hippocampus, for an equal Hebbian response (He et al., 2015; Saylor et al., 2019), which implies that distal predictive neural activity is less persistent under the modulation of the latter (***Figure 1C***). For the navigational task we employed a Morris water maze (MWM) where the agent has to find a hidden platform in a water arena (Vorhees and Williams, 2006;***Figure 1D***). Water is considered a punishing or stress-inducing cue (Harrison et al., 2009), which we hypothesized to cause 5-HT release (Karabeg et al., 2013), and subsequent arrival to the reward zone in the corner of the maze produces an increased dopaminergic response (Frémaux et al., 2013).

**Figure 1:**
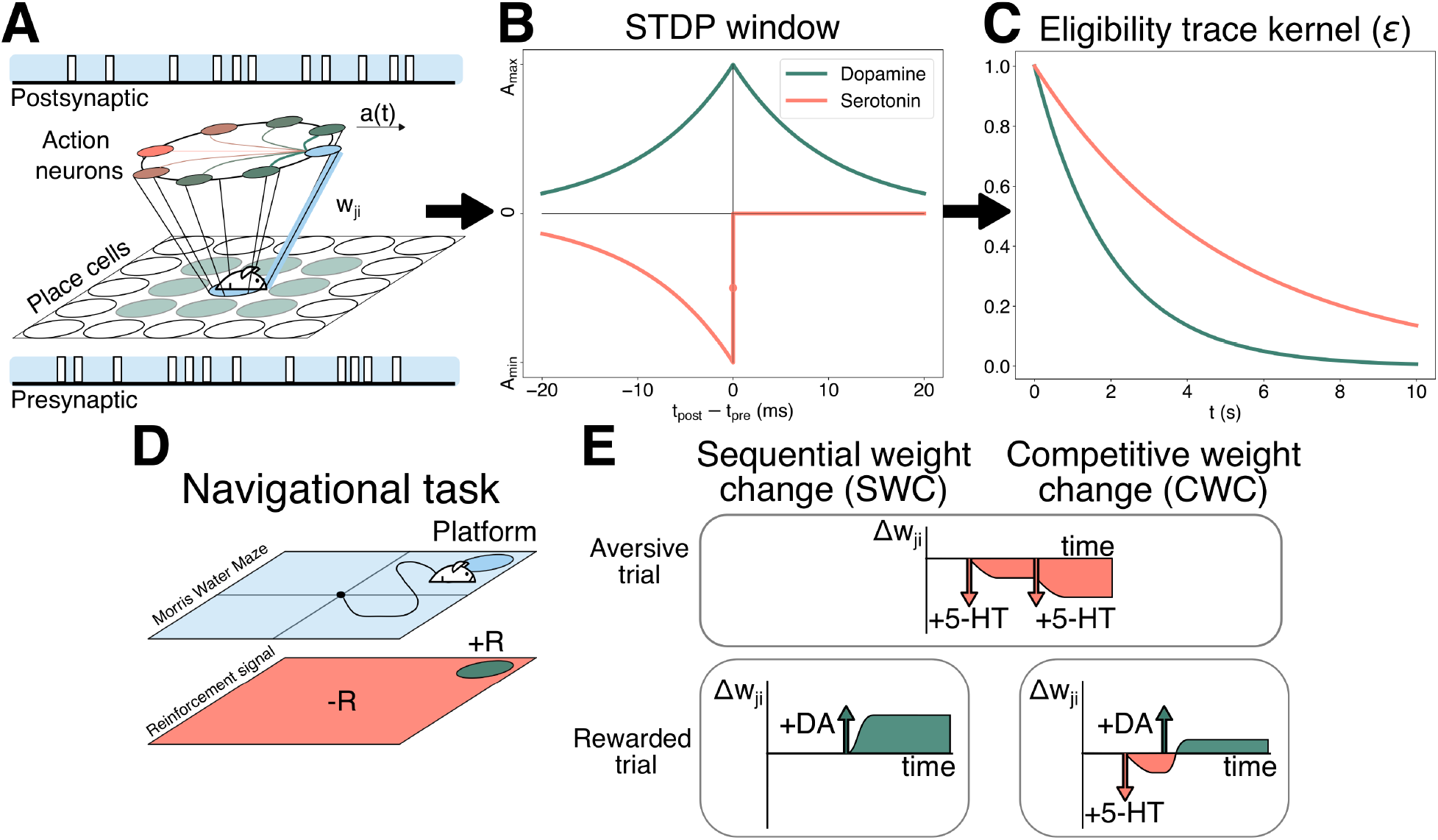
Schematics illustrating the learning model details used in the study. **(A)** The navigational model uses a feed-forward network between Gaussian receptive fields and neurons performing action selection *a*(*t*) through “winner-takes-all” connectivity. Forward synaptic weights *w*_*ji*_ between neuron pairs (blue) are updated through a R-STDP rule. **(B)** STDP window used for DA and 5-HT. Both neuromodulators presented equal decays and opposite modulation with 5-HT presenting exclusively anti-causal depression. **(C)** Kernel of an exponentially decaying eligibility trace with biologically-derived time constants. **(D)** Reward representation derived from water aversion in the Morris water maze (MWM) task. Negative rewards are linked to unsuccessful trials, whereas dopaminergic activation is achieved at platform arrival. **(E)** Strictly aversive trials are processed equally by SWC and CWC. Upon reaching a reward, weight updates for SWC are solely driven by DA. Thus, SWC ensures all connections are either potentiated or depressed at the end of a trial. In CWC, rewarded trials also include aversive cues leading up to the reward. For equations and specific values in **A-C** see Materials and methods.

We considered two R-STDP learning rules to model DA and 5-HT. Firstly, we implemented sequential weight change (SWC), inspired by sequentially neuromodulated plasticity (sn-Plast) (Brzosko et al., 2017; Zannone et al., 2018), in which DA produces t-LTP and 5-HT induces t-LTD during exploration. As sn-Plast, SWC is outcome-dependent and relies on the assumption that either potentiation or depression occurs after a long delay, which allows decoupling rewarding and non-rewarding trials (***Figure 1E*** ; Materials and methods). In the MWM task, SWC produces mutually exclusive t-LTP, when the agent arrives at the reward site, and t-LTD, upon a punishing trial. This is equivalent to consider that DA dominates over 5-HT if the positive valence item is located, which nullifies the contribution of the stress-inducing cue. Conversely, unrewarded trials solely present 5-HT modulation which, in the case of the MWM, is performed at the end-of-trial. Additionally, in opposition to continuous depression (Zannone et al., 2018), we adapted SWC to reflect the presence of an eligibility trace for 5-HT (Materials and methods).

The second learning rule that we examined was competitive weight change (CWC), based on competitive reinforcement learning (Huertas et al., 2016; ***Figure 1D***). In contrast to SWC, eligibility traces perform in-trial opposition (***Figure S2***), defined mathematically as an addition of DA and 5-HT contributions (He et al., 2015; Materials and methods), in which the weight change is determined by the balance between reward-trace pairs. Hence, activated synaptic weights are not guaranteed to be either potentiated or depressed after the reinforcement signal is introduced (***Figure S3***). For some configurations, CWC may yield depressive effects upon reaching the platform if DA does not counterbalance 5-HT. Experimental data from serotonergic and dopaminergic neurons suggests differential activity, since 5-HT is tonically released as a reponse to a long-term punishment, and DA has a greater phasic response towards a rewarding event (Boureau and Dayan, 2011; Cohen et al., 2015). Hence, we represented transient 5-HT as a step function active until the positive valence item is found, which in turn triggers a one-second constant DA signal (Cohen et al., 2015). Consequently, for this navigational task, successful trials include both modulation by DA and 5-HT, whereas unrewarded ones involves exclusively the latter. Fundamentally, SWC and CWC diverge in the timing of the weight update, either end-of-trial or in-trial, and in the characterization of DA and 5-HT as alternate or additive.

Both models were systematically parametrized through grid search to optimize performance. SWC had a better efficiency than CWC in successful simulations over successive trials (***Figure 2A***) and accumulated successful episodes (***Figure 2B***). In both cases, the addition of 5-HT as a t-LTD inducer improved the rewarding outcomes, especially for CWC, which may worsen its learning efficiency with time for particular configurations (***Figure S4***). The poor performance of DA-only CWC can be explained by the fitting of the reward function and the parameters employed, which cause the saturation of weights from non-predictive paths. Compared with SWC, CWC does not optimize the path distance, as measured by the time to reach the reward (***Figure 2C***). Moreover, CWC increases the median distance to the center with time for both conditions (Gehring et al., 2015;***Figure 2D***). The systematic depression of synapses in central place cells, which causes the agent to move near the edges, can explain the apparent incoherence between the latency time and the median distance of CWC with 5-HT. Such a phenomenon is due to the dynamic competition between traces, which can produce depression in neurons activated early under certain conditions. In terms of exploration, we computed the Kullback–Leibler divergence (KL) between the first reward distribution of both conditions (Zannone et al., 2018). The divergence in CWC (KL(+5-HT|−5-HT) = 0.021) and SWC (KL(+5-HT|−5-HT) = 0.015) indicates exploration remains unaltered between both cases (***Figure S5***). For SWC, we also considered a constant action of the punishment, instead of a phasic rewarding response at the end of the trial, without improvements in performance (***Figure S6***). Likewise, in CWC, the efficiency was lowered when a complete phasic response was introduced (***Figure S7***). In conclusion, although limited by dynamical and encoding assumptions, SWC with 5-HT has a better performance than CWC and a dopamine-only learning rule.

**Figure 2:**
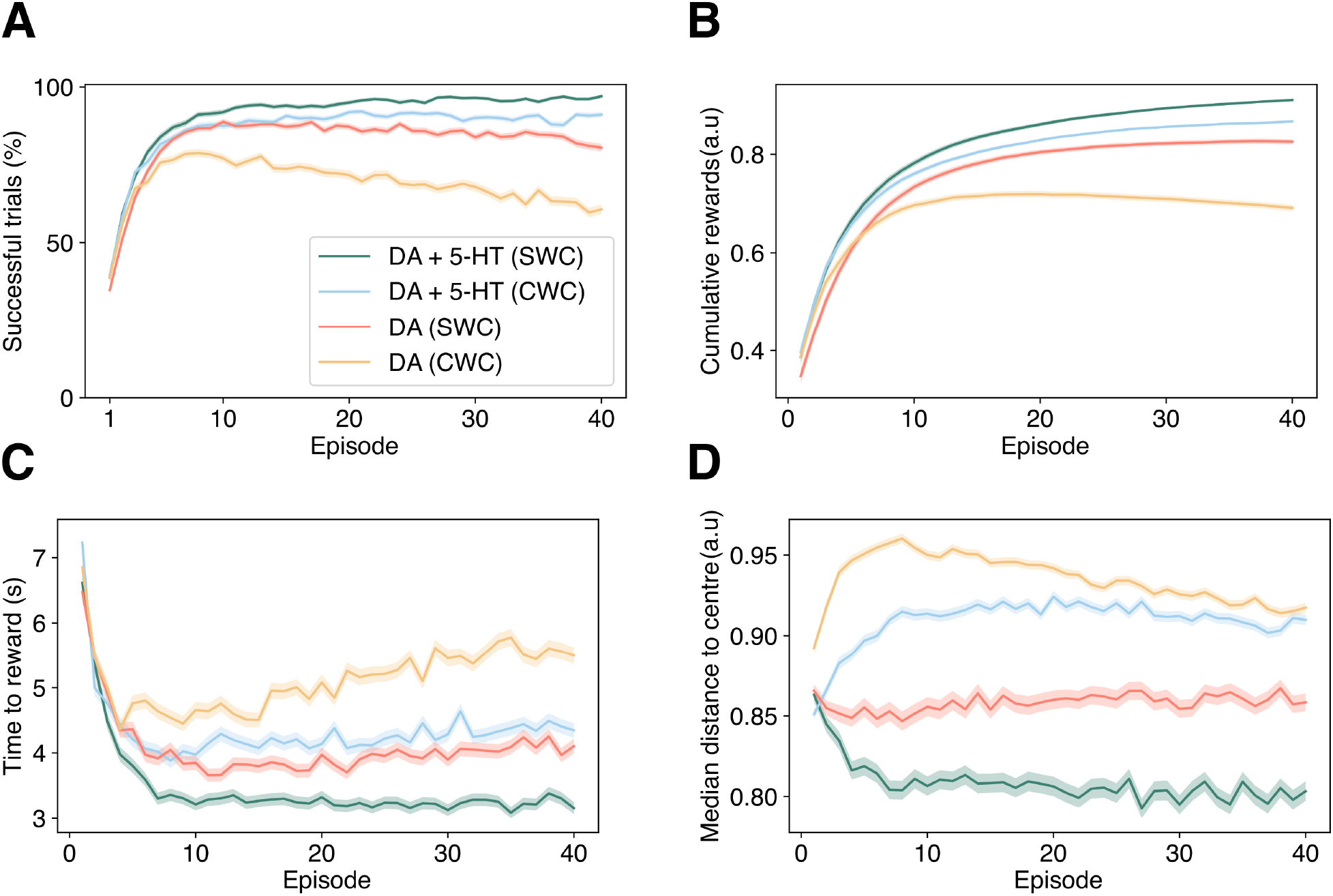
Inclusion of 5-HT improves learning under both R-STDP rules, with enhanced learning for SWC. **(A)** Learning curve of the percentage of successful simulations in each trial. The differences in means between the control (DA) and the addition of 5-HT (DA+5-HT)are significant on the last trial (*p* < 0.01, two-tailed Student’s t-test). **(B)** Cumulative relative number of successful trials averaged over the simulations. **(C)** Average latency time to the reward in positive valence trials. Changes in the time to the reward for the final trial are significant between the different conditions in each rule (*p* < 0.01, two-tailed Student’s t-test). **(D)** Average median distance to the center as measured in spatial memory tests. The shaded ranges correspond to the standard error of the mean (SEM) in M=1000 simulations. See Materials and methods for the value of parameters.

The biological viability of DA and 5-HT modeling through SWC and CWC was assessed against a study by Teixeira et al. (2018), which showed that optogenetic inhibition and activation of serotonergic neurons modified learning abilities of mice without significantly affecting locomotion. We aimed at replicating these results by imposing three intervals of the simulation time increasing or omitting the serotonergic signal (***Figure 3A***), considered as a doubling and absence of the punishment, correspondingly. Notably, inhibition of 5-HT in CWC caused a performance reduction although overactivation yielded no variation (***Figure 3A.i***). In contrast, SWC worsened the number of rewarding trials under either condition (***Figure 3A.ii***).

**Figure 3:**
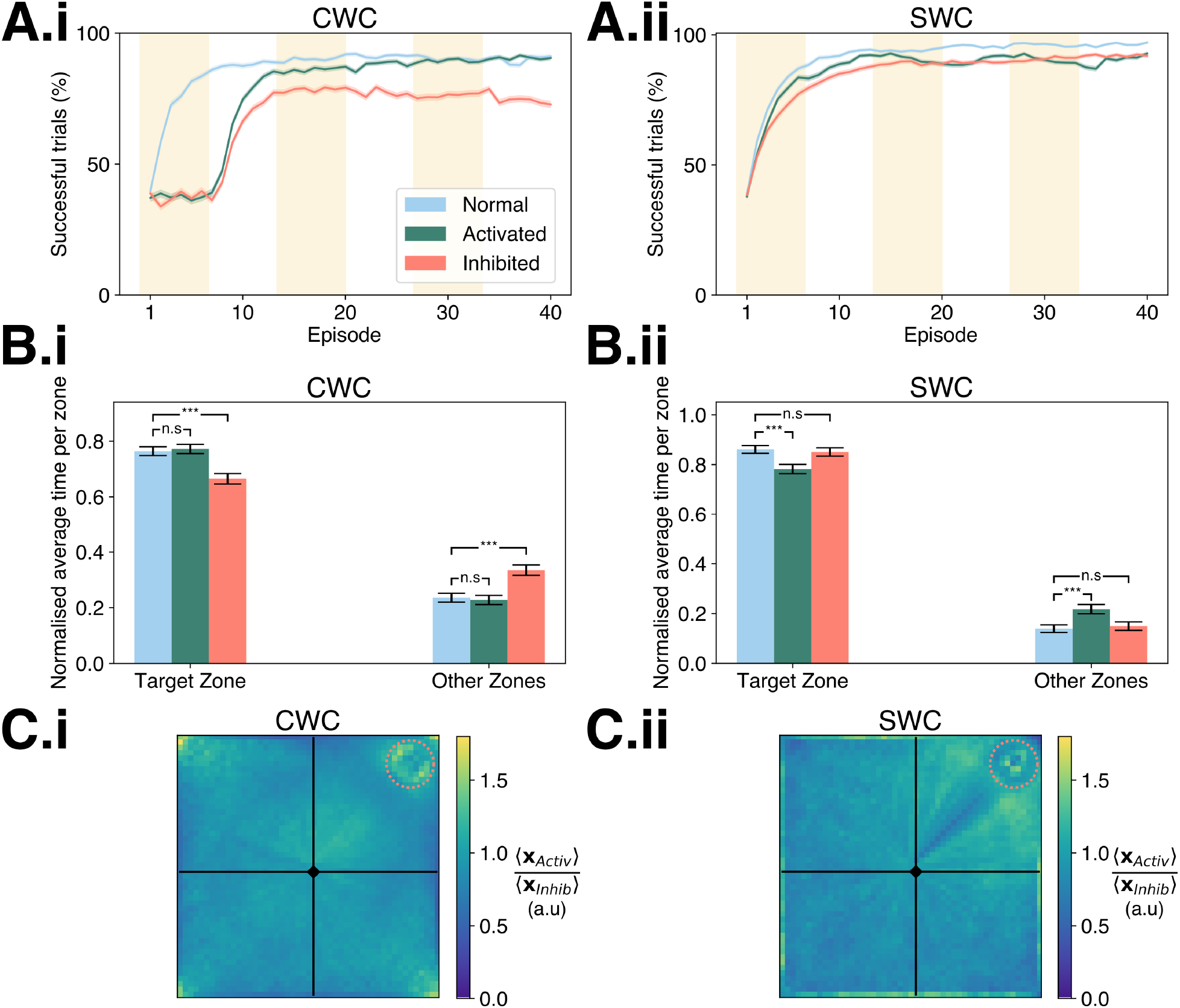
Overactivation and inhibition of 5-HT in CWC resembles behavioral data from optogenetic modulation of serotonergic neurons in the MWM. **(A)** Average percentage of successful simulations for (i) CWC and (ii) SWC. Episodes in yellow correspond to times in which optogenetic changes are introduced. **(B)** Bar plot of the average time in the target quadrant against the other zones for (i) CWC, which resembles real data observed in mice from Teixeira et al. (2018) and (ii) SWC. Statistical significance (two-sample Student’s t-test with *p* < 0.001, ***) is shown for changes in some conditions under paired test . **(C)** Fold change between activated and inhibited averaged over time and simulation position histograms (50 bins per side) for **(i)** CWC and **(ii)** SWC. In a trial, the position is binned spatially then averaged across episodes and trials. The ratio of mean location between conditions is shown with the initial position (rhombus) and the reward location (dotted circle). Filled area and error bars in **A-B** correspond to SEM (M=1000).

These results highlight the importance of 5-HT as a compensatory mechanism of DA in CWC against a greater role in negative sampling for SWC. For both learning rules, sequential 5-HT inhibition decreased the residence time at the target platform in contrast with the control (***Figure 3B***), albeit only significantly for CWC (***Figure 3B.i***). Nevertheless, the increase in serotonergic response only resulted in a larger amount of time spent at the target zone for CWC (***Figures 3B.i-ii***). Hence, in time spent in the target quadrant, CWC replicated the changes observed empirically but SWC did not. A comparison between the position traces for activation and inhibition shows that 5-HT intensifies movement near the starting position and the corners of the maze in CWC (***Figure 3C.i***) and a more systematic navigation of the target quadrant in SWC (***Figure 3C.ii***), in which serotonergic amplification impedes the learning of the shortest path. Overall, these results suggests a greater biological feasibility of CWC in DA and 5-HT modeling. Nonetheless, additional evidence regarding the activity of serotonergic and dopaminergic neurons during individual trials could provide a more robust test for the model and corroborate the LTD contribution of 5-HT.

The described valence system has been evaluated in reversal learning, which involves punishment and reward switching (Matias et al., 2017). In this setting, t-LTP and t-LTD are disjointed (i.e., the agent is either rewarded or punished in the same trial), which reconciles SWC and CWC rules into the same system with phasic activity (***Figure S8***). As 5-HT operates under different timescales in reversal (Matias et al., 2017), we evaluated five learning rates to quantify the relative importance of t-LTD in relearning. For all rates, forward learning was successful (***Figure 4A***) although there was no recovery after inversion (***Figure 4B***), with high learning rates performing better. The lack of discrimination is observable in post-reversal synaptic weights compared to forward ones (***Figure 4C***). Additionally, negative valence simulations decreased for all learning rates (***Figure 4D***) although most simulations were neutral (***Figures 4E***), indicating the emergence of a non-decisive or metastable state in deliberation (Bakkour et al., 2019). Reduced selectivity is explained by low polarization in feed-forward weights, as measured by the coefficient of variation (***Figure 4F***), for all learning rates. Taken together, these results predict a role of 5-HT in aiding reversal learning and present testable conditions in open field navigation under rewarding and punishing cues. However, this model does not explain positive reward encoding of 5-HT observed in conditioning trials (Matias et al., 2017), which suggests a more complex interplay between neuromodulators in valence learning.

**Figure 4:**
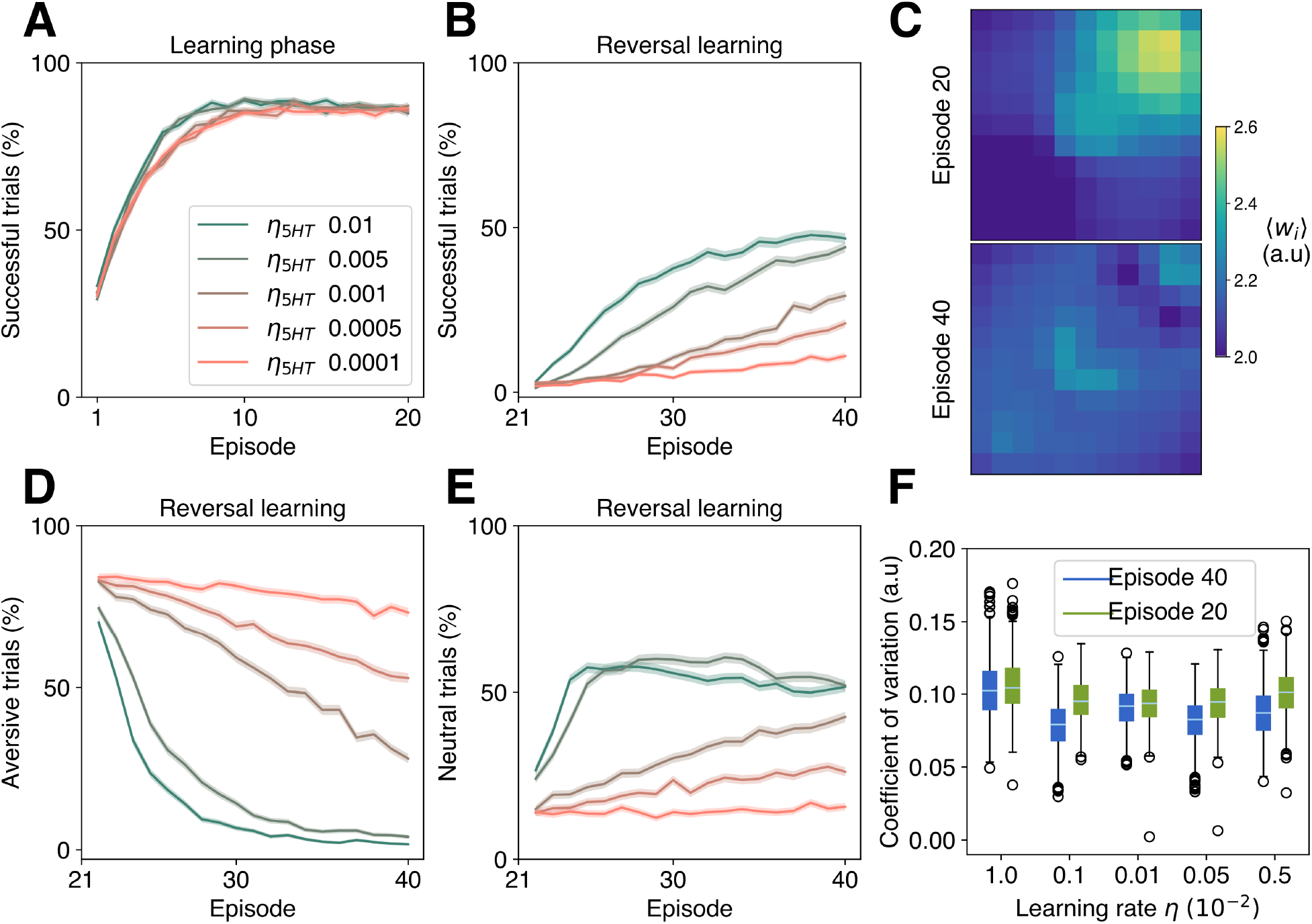
Reversal learning in an open field improves under 5-HT modulation for different learning rates with a high preference for a neutral state. **(A)** Learning curve depicted as the average percentage of successful simulations per trial for five 5-HT learning rates before reversal. **(B)** Average successful simulations after reversal. **(C)** Average place cell weights before (trial 20) and after reversal (trial 40) for *η*_5*HT*_ = 0.01. **(D)** Average punishing or negative simulations after reversal. **(E)** Average latency time to the reward in successful simulations. **(F)** Distribution of the mean coefficient of variation (CV) of synaptic weights before and after inversion. This corresponds to the ratio of the standard deviation to the mean of weights in each place cell. Lower CV values after inversion imply a decrease in dispersion or polarization of synaptic weights, as shown in **(C)**. In curves, the shaded region corresponds to SEM (M=1000).

In summary, our plasticity model of interacting DA and 5-HT contributions provides mechanistic insights into their role in hippocampal-dependent spatial navigation. Future extensions of this model could explore combinations of aversive and attractive states coded at the level of circuits.

## 3 Materials and methods

Hippocampal-dependent spatial navigation is modeled through a one-layer network based on a navigational actor-critic system (Frémaux et al., 2013; Brzosko et al., 2017). Location, encoded through the spiking rate of place cells, serves as the input. The output layer is composed of action neurons, which determine the preferred movement of the agent by their firing rate.

### 3.1 Place cells

The spiking activity of place cells represents two-dimensional positional information. These are modeled through an inhomogeneous Poisson process with maximum spiking activity 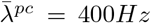. The squared norm of the deviation between the Cartesian location of the agent **x**(*t*) and the center of the place cell *i* is used for the calculation of the rate as follows:

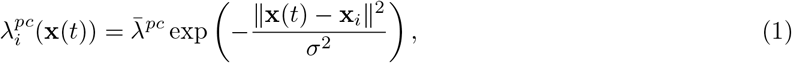

The firing rate of the Poisson process exponentially decays with the Euclidean distance between the agent and the center of the place cell. A total of 121 place cells, equally separated by a distance of *σ* = 0.4 a.u, were distributed on a square of length side 4 a.u (Brzosko et al., 2017).

### 3.2 Action neurons

To model action neurons, a zero-order Spike Response Model (*SRM*_0_) was used (Gerstner, 1995), in which the membrane potential is represented as

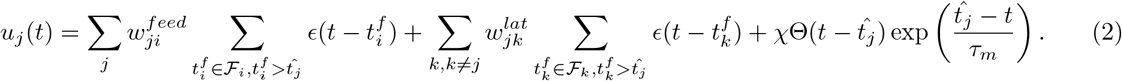

In the feed-forward network, the action neuron *j* receives an excitatory postsynaptic potential (EPSP) 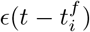, from place cell *i*, for firing times in the set 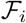, after the last spike of action neuron 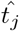 and under a synaptic efficiency 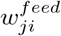. Similarly, action neurons *k*, of the lateral connectivity network, are with synaptic weights 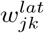 connected and their spike arrival times are contained in set 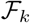. The EPSP kernel is

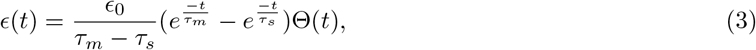

where Θ(*t*) is the Heaviside step function, the membrane constant *τ*_*m*_ = 20 ms and the rising time *τ*_*s*_ = 5 ms. Eq. 2 considers a scale factor for the refractory effect *χ* = −5mV and Eq. 3 as well with *∊*_0_ = 20 mV·ms.

The spiking activity is determined by an inhomogeneous Poisson process with rate *λ*_*j*_(*u*_*j*_(*t*)), which is formulated as

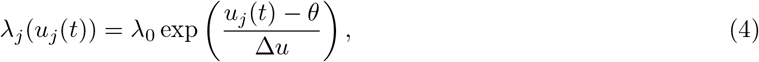

in which the maximum rate is *λ*_0_ = 60 Hz, the potential threshold is *θ* = 16 mV and Δ*u* is a voltage window for spike emission that determines the degree of randomness. The dynamics are simplified if the resting potential is assumed to be 0V (Zannone et al., 2018).

The instantaneous firing rate of an action neuron *ρ*_*j*_(*t*) is obtained by filtering the spiking activity 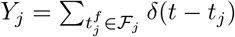 with the kernel 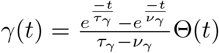, where *τ*_*γ*_ = 50ms and *ν*_*γ*_ = 20 ms.

Each action neuron *k* represents a preferred direction of movement **a**_*k*_ and interacts with other angle-encoding action neurons through a lateral connectivity (Frémaux et al., 2013). The lateral synaptic weight dynamics produce a “N-winner-takes-all” arrangement by which *N*_*action*_ neurons compete for the preferential angle. Hence, the connectivity between neural units *k* and *k′* is modeled to inhibit opposite directions and excite similar ones as

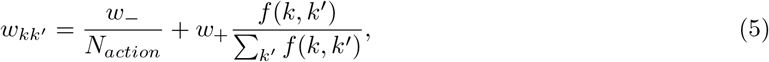

inhibitory and excitatory weights are *w*_−_ = −300 and *w*_+_ = 100 and *f* is the lateral connectivity function, which reaches a maximum for *k* = *k′* ± 1 and decreases monotonically towards zero for *k* = *k′*. Concretely, in this case, *f*(*k*; *k′*) = (1 − *δ*_*k,k′*_) exp(*ζ* cos(*θ*_*k*_ − *θ*_*k′*_)) decreases exponentially for increasingly dissimilar angles *θ*_*k′*_ and is scaled by a factor *ζ* = 20. These parameters were tuned for a population of *N*_*action*_ = 40 with 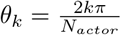 (Zannone et al., 2018). Thus, action vectors were **a**_*k*_ = *a*_0_(sin(*θ*_*k*_), cos(*θ*_*k*_))^*T*^ with *a*_0_ = 0.08.

The action resulting from the spiking activity of the network is coded through a population vector as

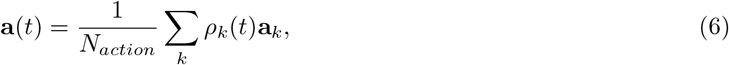

which is weighted by the filtered spiking activity of neurons *ρ*_*k*_(*t*) = (*Y*_k_ ○ *γ*)(*t*). Thus, the action at each time step **a**(*t*) is computed as the average of the action vectors with the predicted instantaneous activity of actor neurons. The inertia of movement is determined by the activity of the network with the maximum velocity being limited by *a*_0_.

### 3.3 Navigational setting

The square *S* delimiting the two-dimensional plane, serves as a boundary condition for the position **x**(*t*) and the movement of the agent **a**(*t*). This is formulated as

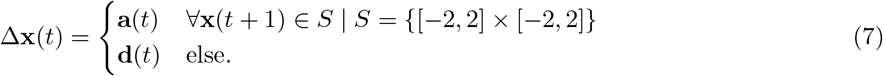

The bouncing vector **d**(*t*) corresponds to the displacement of the agent in the direction of the normal vector to *S* or **n**_*S*_, which is defined as **d**(*t*) = *d*_0_**n**_*S*_(**x**(*t*)). The bouncing distance is set as *d*_0_ = 0.01.

In the Morris water maze, the rewarding platform was positioned at *r*_*c*_ = (1.5, 1.5) with a radius *r*_*r*_ = 0.3. In reversal learning, the reward is initially maintained at its position and the punishment is placed at *p*_*c*_ = (−1.5, −1.5) with radius *p*_*r*_ = 0.3, for trials 1 to 20. After reversal, both elements switch position, *r*_*c*_ = (−1.5, −1.5) and *p*_*c*_ = (1.5, 1.5), with unvaried radii, for trials 21 to 40. In all instances, the initial position of the agent was **x**(*t* = 0) = (0, 0) and the maximum trial time was *T*_*max*_ = 15 s (Zannone et al., 2018). If the reward or, in reversal learning, punishment is reached before *T*_*max*_, the trial is ended, and place cells are deactivated. In sequential weight change, the weight update occurs at *t* = *T*_*rew*_ +300 ms, to replicate consummatory behavior. Per contra, in competitive weight change, weights are updated continuously until the DA signal is no longer active *t* = *T*_*rew*_ + *T*_*DA*_. Activity is reset between trials.

### 3.4 Sequential weight change (SWC)

The sequentially neuromodulated plasticity (sn-Plast) rule from Brzosko et al. (2017) was adapted to match the empirical evidence available for 5-HT in He et al. (2015). Hence, instead of presenting an online depression mediated by acetylcholine, the adjusted SWC update introduces an eligibility trace ϵ_5*HT*_ for the depressor, which permits the temporal discrimination of neural activity leading up to an aversive cue.

The weight update is determined by the spike-timing-dependent plasticity (STDP) window of each neuromodulator *W* (*s*), filtered by its eligibility trace kernel *ϵ*, and an outcome-dependent signal *A* that . As such,

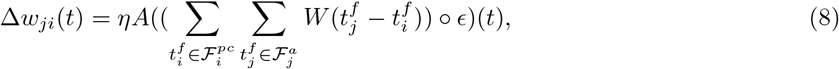

with differentiated learning rates *η* for 5-HT and DA 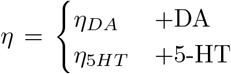. SWC assumes either DA or 5-HT act at the end-of-trial. Thus, in the MWM, DA is released if the agent reaches the reward while, in an unrewarded trial, water aversiveness causes 5-HT stimulation. This representation holds for reversal learning, outcomes are exclusive as well. Hence, the valence signal *A* is conditioned by whether the trial was rewarding or punishing and it is defined as 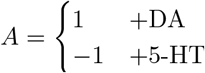. The firing times 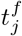 and 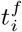 of action neuron *j* and place cell *i* are contained in the sets 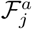 and 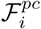.

The STDP window for DA was preserved as 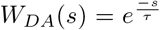, where *τ* = 10 ms. However, for 5-HT (He et al., 2015), it was transformed into an asymmetric STDP curve as

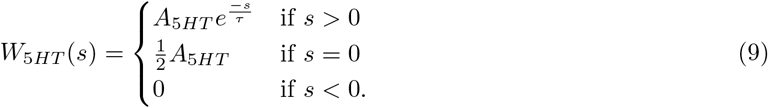

The eligibility trace kernel for 5-HT and DA is formulated as

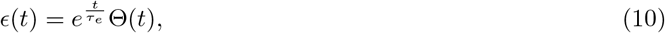

with 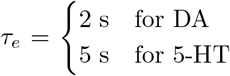. The value of the time constant for 5-HT comes from *in vivo* experiments (He 5 s for 5-HT et al., 2015). The decay disparity among traces elicits a differential response between predictive neural “trails”. In other terms, place cells encoding for the same path suffer a greater weight change, in absolute terms, through serotonergic action than with DA modulation.

### 3.5 Competitive weight change (CWC)

In competitive weight change (CWC), inspired by competitive reinforcement learning (CRL) from Huertas et al. (2016), the eligibility traces, or proto-weights, of the two modulators are under a dynamic competition for the upgrade of the synaptic weight. Proto-weights *T* are the result of filtering the Hebbian term *W* (*s*) for each neuromodulator with the eligibility trace kernels ϵ defined in Eq. 10. The equation results in

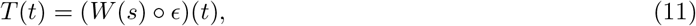

with the STDP windows for DA and 5-HT defined as in SWC In integral terms, and for a particular connection between neurons *j* and *i*, Eq. 11 becomes 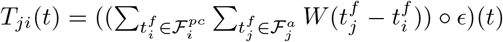.

The weight uptade is expressed as the dynamic competition between traces, which corresponds to the difference between t-LTP and t-LTD proto-weights or

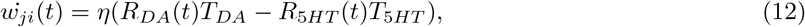

where *T*_*DA*_ and *T*_5*HT*_ are the proto-weights of the neurotransmitters and *R*_*DA*_ and *R*_5*HT*_ models the reinforcement response for each neuromodulator with learning rate *η*. Hence, the condition for t-LTP or t-LTD becomes

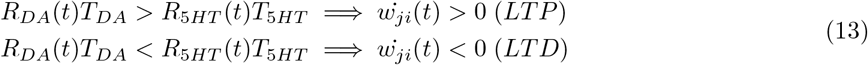

The integral version of Eq. 12 is,

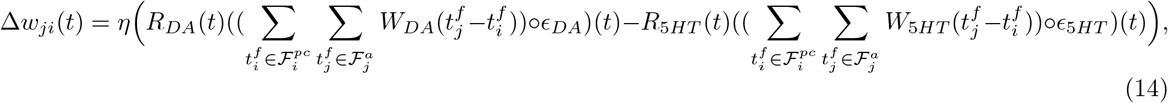

where the reinforcement signals are modeled as Heaviside step functions Θ(*t*). Specifically, *R*_*DA*_(*t*) = *R*_*DA*_(Θ(*t − T*_*rew*_) − Θ(*t − T*_*DA*_ − *T*_*rew*_)) and *R*_5*HT*_ (*t*) = *R*_5*TH*_ (Θ(*t*) − Θ(*t − T*_*rew*_)). Accordingly, 5-HT is active until the reward is attained or the end of the trial at time, if the agents has been unsuccessful, which corresponds to time *T*_*rew*_. DA acts for a time after the arrival of the agent to the positive reinforcement site, which was assumed to be *T*_*DA*_ = 1 s in accordance with biological studies involving long-term observation of the dopamine-serotonin interplay (Cohen et al., 2015).

The definition from Eq. 12 In opposition to the formulation by Huertas et al. (2016), the eligibility traces of LTP and LTD were not such that could be used for the representation of cues through their dynamic equilibrium. Alternatively, at the end of the simulation, the potentiation or depression of flagged synapses (i.e., neural connections that have been active during the trial) is not assured and may not be consistent with the outcome of the trial. Notably, in our MWM implementation, the condition for weight potentiation after reaching the platform requires that

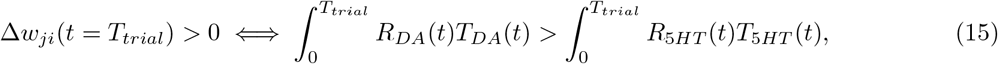

in which *T* is the proto-weight of each neuromodulator and *R* the reinforcement signal. It should be noted that this equation depends on the time of the trial *T*_*trial*_ and, therefore, the sign of weight update values can vary between trials of different duration. This is equivalent, in integral terms, to Eq. 13.

### 3.6 Parameter values

The configuration of each model was optimized through grid search parametrization. In particular, we optimized the amplitudes of the STDP widows, *A*_*DA*_ and *A*_5*HT*_, the reward magnitudes for CWC *R*_*DA*_ and *R*_5*HT*_, and the learning rates *η*. Sweeps of these values were first conducted by orders of magnitude and then with fine tuning around good estimates. The best models were selected by the proportion of successful simulations at the final trial.

**Table 1:** Parameters used for SWC and CWC models. These parameters correspond to the best models with regard to the percentage of successful simulations in the last trial.

| Model | $A_{DA}$ | $A_{5HT}$ | $\eta$ | $w_{min}$ | $w_{max}$ | $R_{DA}$ | $R_{5HT}$ | $\tau_{STDP}$ | $\tau_{\epsilon}(DA)$ | $\tau_{\epsilon}(5HT)$ |
| --- | --- | --- | --- | --- | --- | --- | --- | --- | --- | --- |
| CWC | 1 | 0.01 | 0.0001 | 1 | 3 | 1 | 1 | 10 ms | 2 ms | 5 ms |
| SWC | 1 | 1 | 0.01 | 1 | 3 | - | - | 10 ms | 2 ms | 5 ms |

The code, written in Python, will be available after publication in ModelDB.

## 4 Acknowledgments

This work was funded by BBSRC (BB/N013956/1 and BB/N019008/1), EPSRC (EP/R035806/1), German Research Foundation (CRC 1089), “la Caixa” Foundation (LCF/BQ/EU19/11710071), Max Planck Society, Simons Foundation (564408) and Wellcome Trust (200790/Z/16/Z).

## Supplemental Information

**Figure S1:**
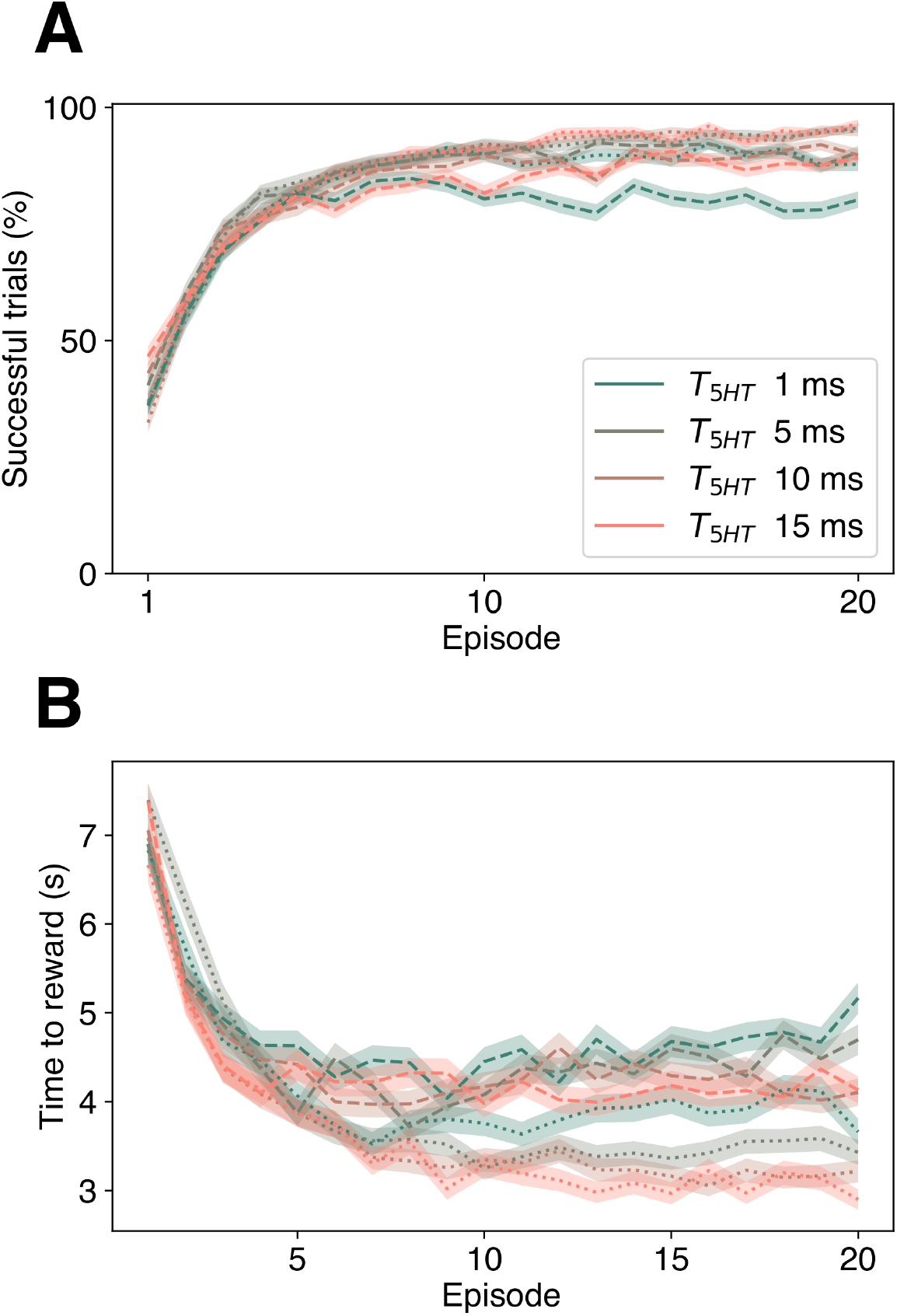
STDP decay value of 5-HT does not hinder learning. **(A)** Learning curve depicted as the percentage of successful trials along episodes. Neither SWC (dotted) nor CWC (dashed) altered significantly their performance for different decays of the exponential kernel of the STDP window (Eq. 9, Materials and methods). **(B)** Latency time. Greater variation can be seen among the times to reach the reward and shortest path optimization. Higher time constants for 5-HT-dependent STDP facilitate greater distance minimization. Filled areas correspond to SEM (M=1000).

**Figure S2:**
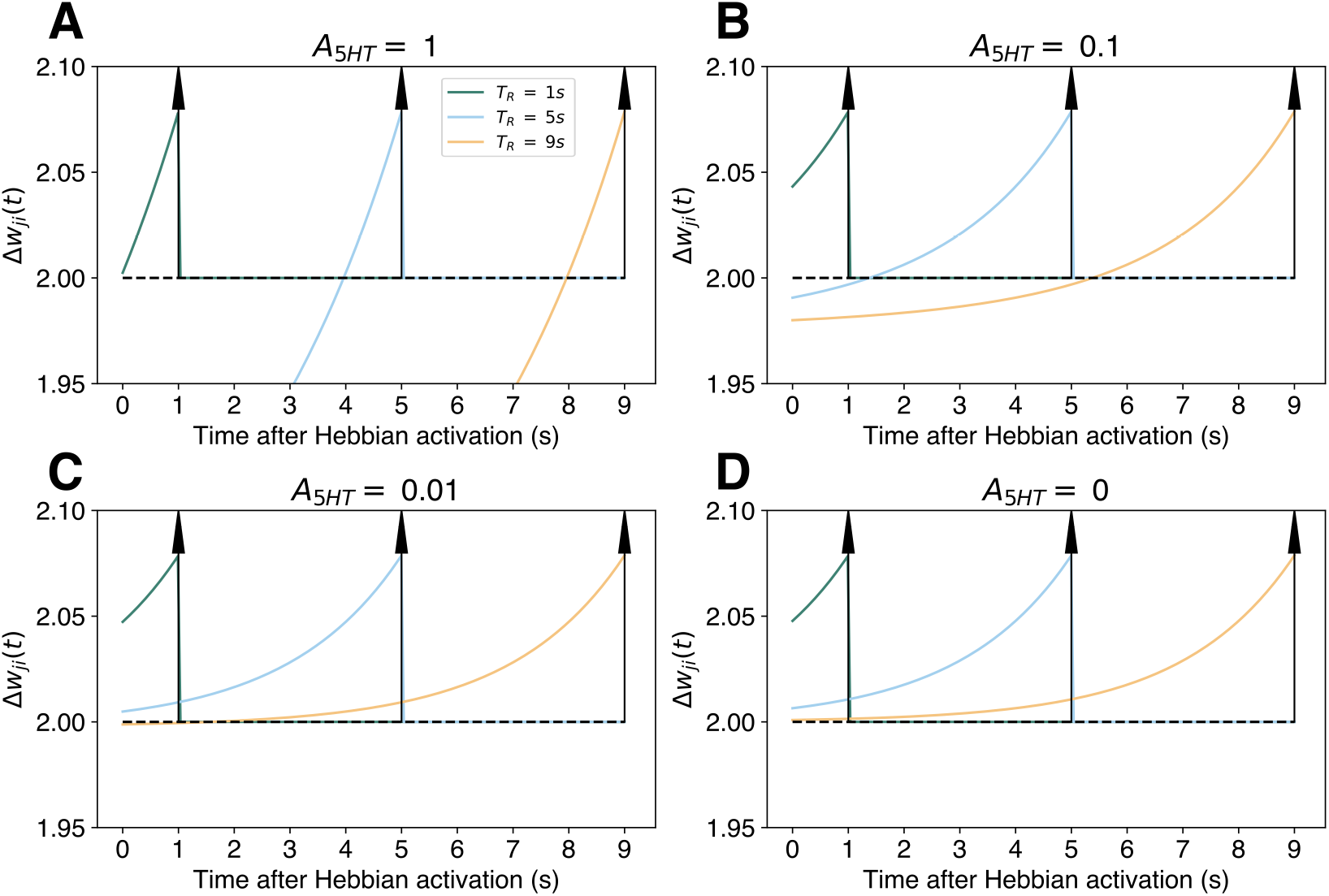
SWC performs sign-consistent updates across all weights. Weight change as a function of the time after a Hebbian activation 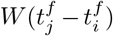 (Eq. 8), modeled as a Dirac delta function at *t* = 0 or *δ*(*t*). Three different rewarding times *T*_*R*_, at 1s, 5s and 9s, and a punishment at the end of the episode are shown (arrows). With the weight update Δ*w*_*ji*_(*t*) for each time after Hebbian activation plotted. The parameters were taken as specified in Materials and methods (Table 1) with the variation of the magnitude of the STDP window of 5-HT. Four different values **A-D** for serotonergic modulation are used (i.e. *H*_5*HT*_ (*t*) = *A*_5*HT*_ δ(*t* − *t*_*activation*_).

**Figure S3:**
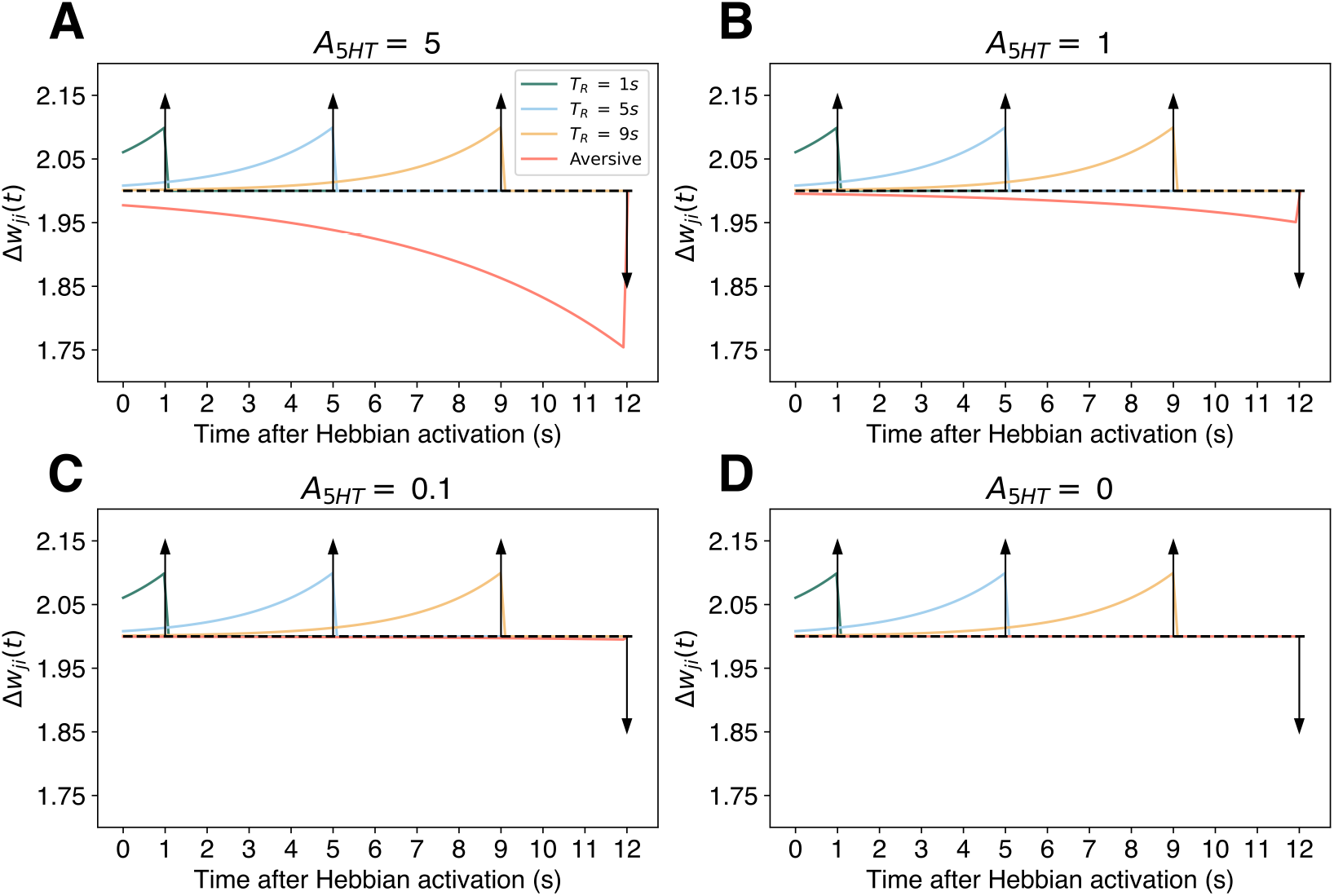
Potentiation of positive valence trials in CWC is not guaranteed for all neurons leading up to the reward. This case represents CWC for serotonin and dopamine with a Dirac delta function at *t* = 0 or *δ*(*t*) as the Hebbian term. Three rewarding time points *T*_*R*_ are shown (vertical arrows), which denote the time at which serotonergic and dopaminergic responses are switched (i.e. *R*_*DA*_(*t*) = *δ*(*t* − *T*_*R*_)). The weight update Δ*w*_*ji*_(*t*) is plotted against the time after the Hebbian activation. As described (Materials and methods), We maintained dopaminergic response for 1s as consummatory behavior. Four different values (**A-D**) for serotonergic modulation are used (i.e. *H*_5*HT*_ (*t*) = *A*_5*HT*_ δ(*t* − *t*_*activation*_). Depression occurs for all ranges of modulation (**A-C**) except for *A*_5*HT*_ = 0 (**D**).

**Figure S4:**
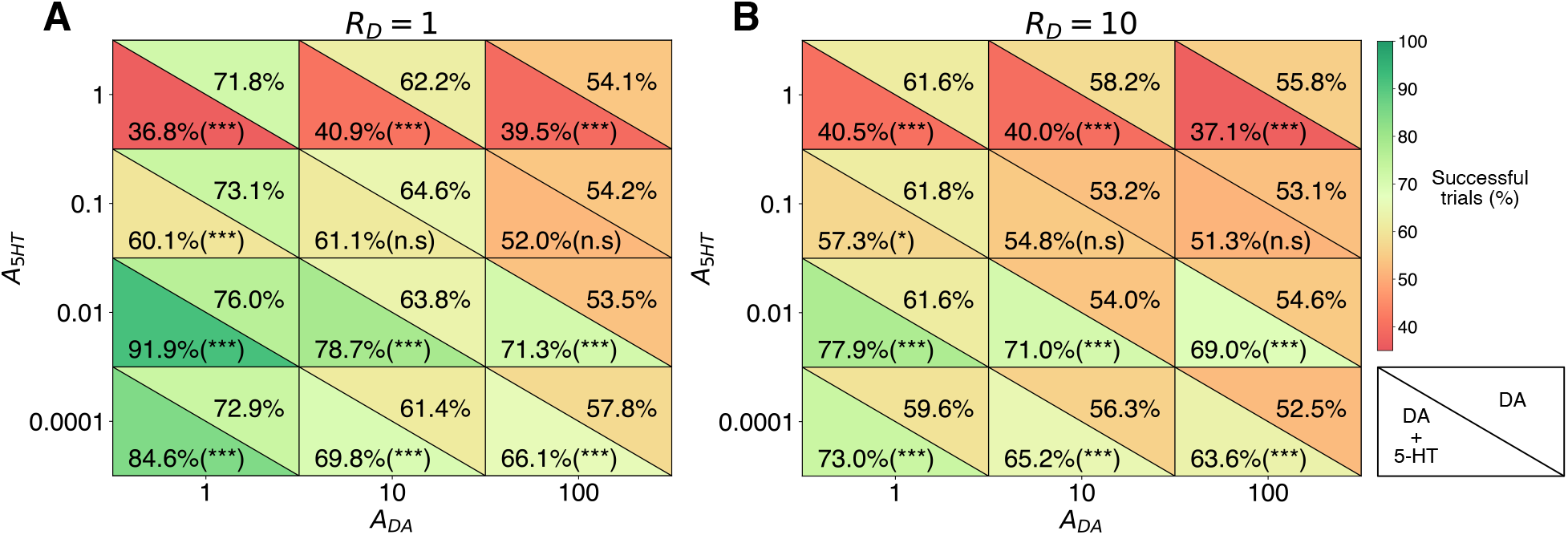
Manual search of the amplitudes of the STDP window for DA and 5-HT. Scores at each amplitude tuple correspond to the mean percentage of successful trials at episode 20 (M=1000) for runs with DA and 5-HT (bottom left) and DA-only (upper right). Two values of 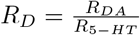 were used (Eq. 14, Materials and methods). Equal reward functions in strength (**A**) achieved better scores than a ten-fold difference (**B**). For some configurations, DA-only performs better than the competition between neuromodulators. However, the highest efficiencies in both tables correspond to a joint regulation. Changes in success rates between conditions were tested for statistical significance (two-sample Student’s t-test with *p* < 0.001, ***)

**Figure S5:**
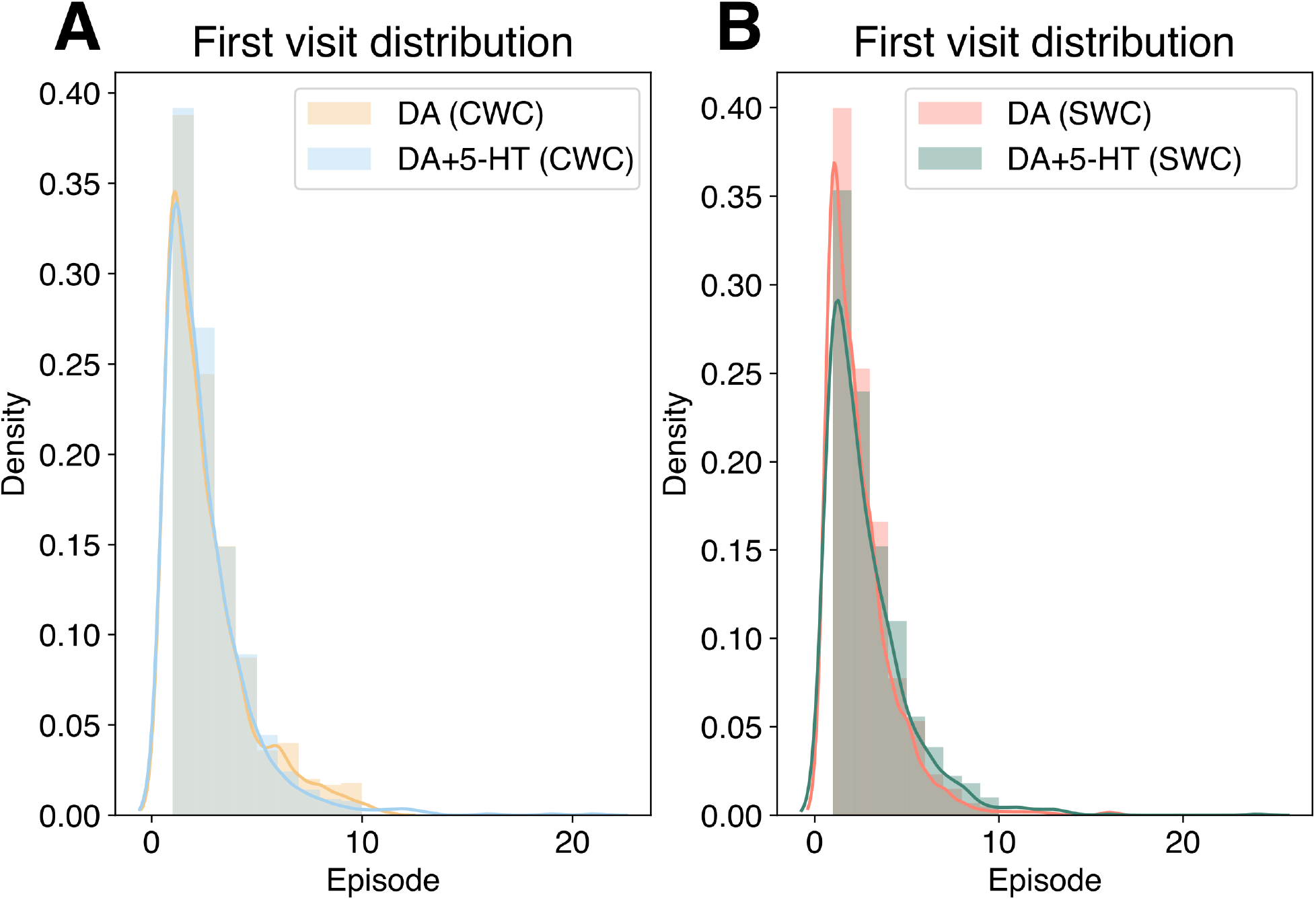
Addition of 5-HT does not alter explorative behavior, depicted as first visit distribution. **(A)** Comparison of the histograms of the first reward visit for 5-HT+DA and DA only in CWC (KL= 0.021). **(B)** Difference between the sample distribution of the first reward visit for 5-HT+DA and DA only in SWC (KL=0.015). Lines correspond to smoothing with kernel density estimation. Each conditioned was sampled for M=1000 simulations.

**Figure S6:**
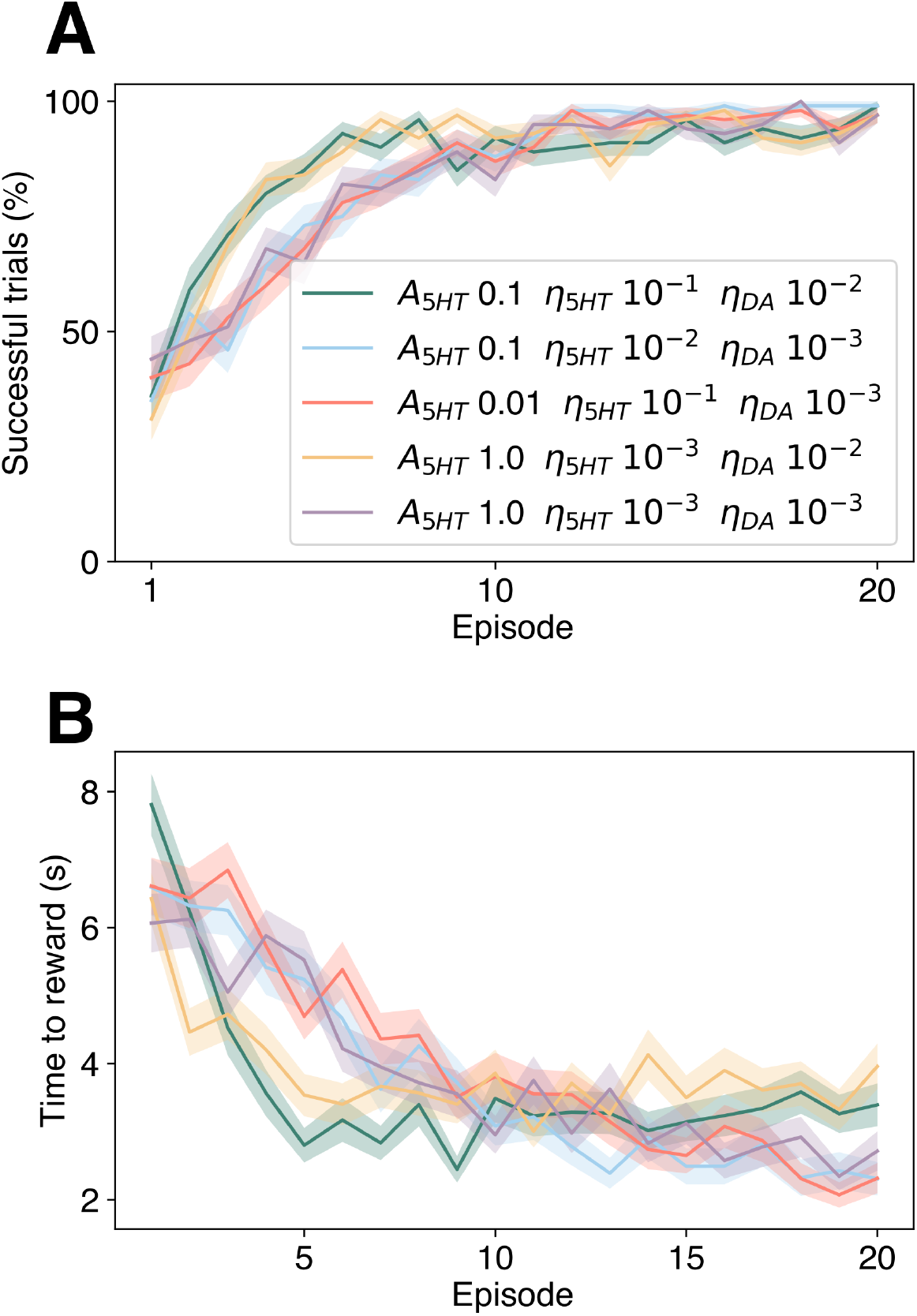
A step response of 5-HT in SWC does not impede learning in a MWM task. We tested a Heaviside function normalized to have the same maximum total response in a trial as the delta function formulation (i.e. 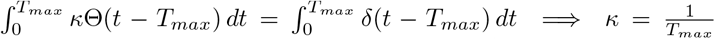). **(A)** Learning curve of the best four STDP window combinations of nine tested. **(B)** Latency time for the best four Hebbian modulation formulations. In both cases, the average is shown with the filled area corresponding to the SEM (M=100 simulations). All other parameters were maintained constants.

**Figure S7:**
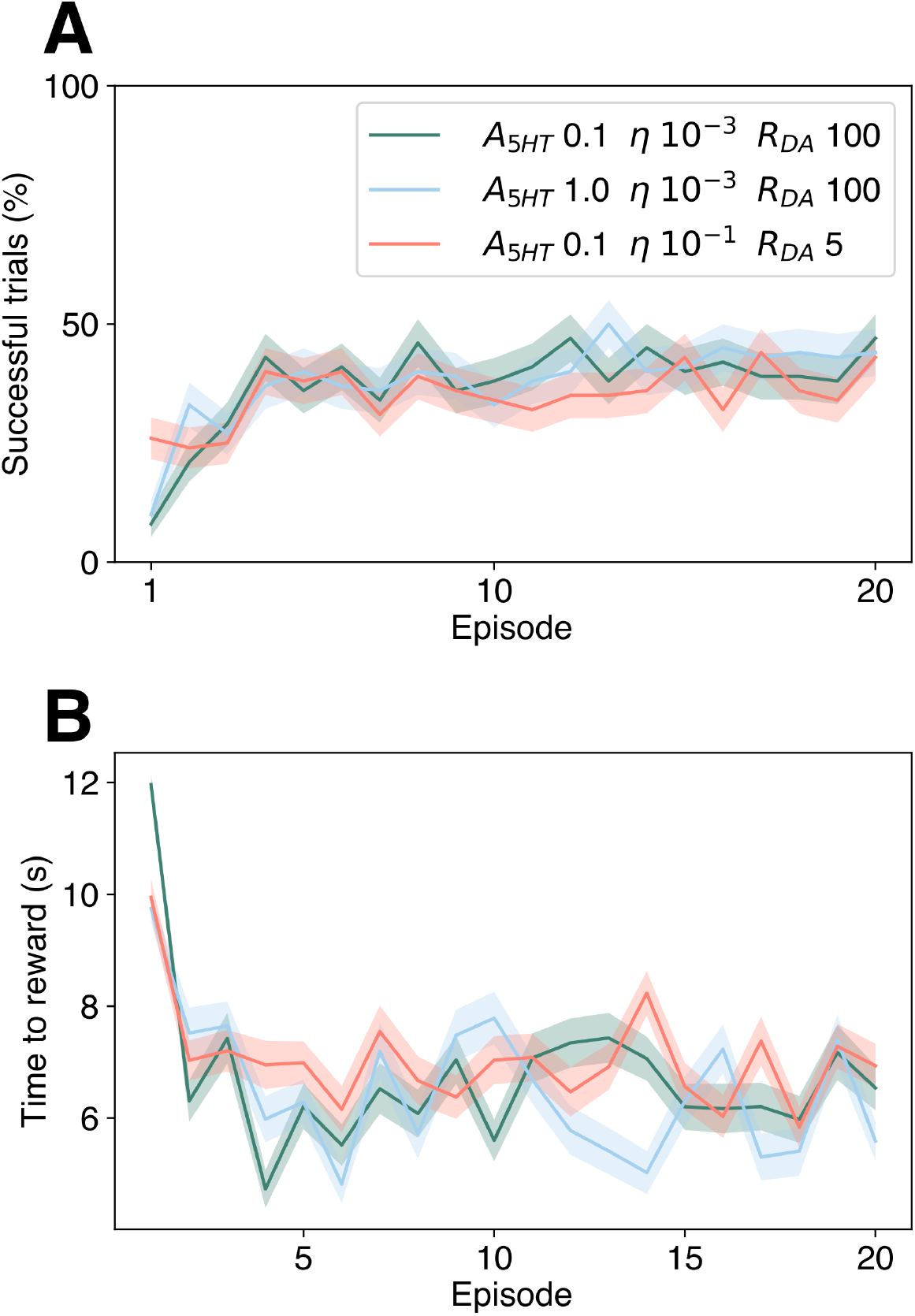
Dirac delta DA response hinders learning performance. We employed a Delta function formulation at the time of the reward with a posterior 300 ms continuous weight update. Thirty models were trained for different combinations of reward strength, STDP window of serotonin and learning rates. **(A)** Learning curve of the best three combinations in the percentage of successful trials. **(B)** Latency time for the best three configurations. In both cases, the line corresponds to average per test, and the shaded area corresponds to SEM (M=100 simulations).

**Figure S8:**
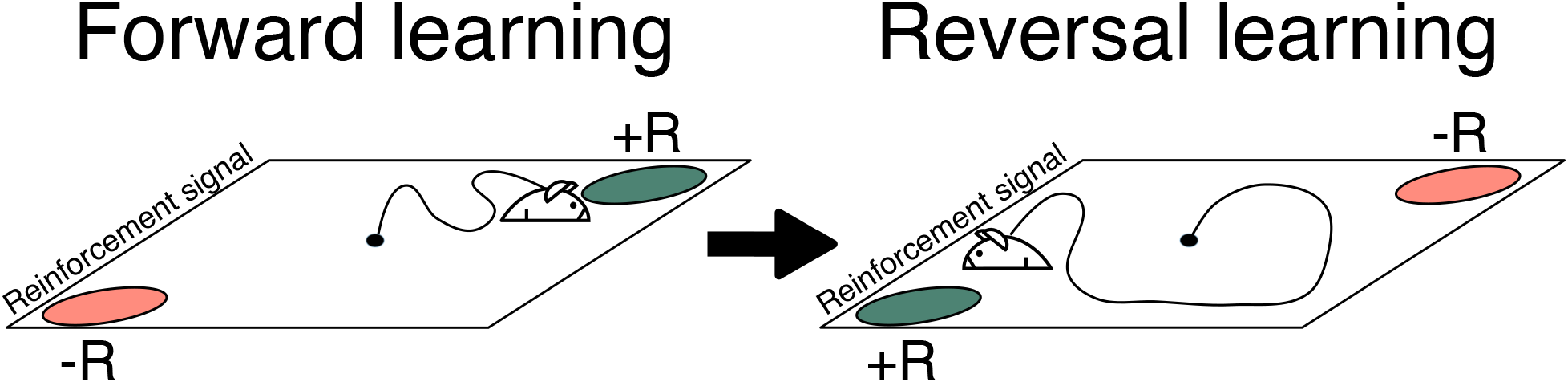
The task in reversal learning consists of an open-field (same characteristics as in Eq. 7; Materials and methods) with a reward (upper right) and a punishment (lower left). We assumed that the reward and the punishment are modeled with Dirac delta functions at the time the reinforcement signal is active *T*_*rew*_ (i.e. *R*(*t*) = *Rδ*(*t* − *T*_*rew*_)), with 300 ms of consummatory time for each neuromodulator (Materials and methods). If none of the items is reached in *T*_*max*_ the trial ends. In discrete form, the CWC rule transforms to

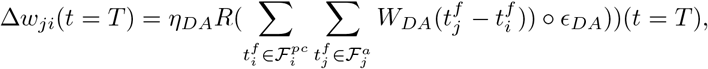

which, with separated learning rates for each neurotransmitter, is identical to presenting different STDP window amplitudes. Thus, it is equivalent to SWC in this setting.

